# Mapping the Structural Landscape of Amyloid Fibrils to Guide Polymorph-Specific Therapeutics

**DOI:** 10.1101/2025.05.08.652887

**Authors:** Ahmed Sadek, Bruno E. Correia, Hilal A. Lashuel

## Abstract

Amyloid fibrils are pathological hallmarks of neurodegenerative diseases and central contributors to their progression, representing promising targets for disease-modifying interventions. However, limited access to patient-derived fibrils and the inability to reproduce pathological folds *in vitro* hinder the development of fibril-specific ligands. Here, we present FibrilSite, a computational pipeline that identifies geometric and physicochemical similarities across fibril surfaces and demonstrate its utility to identify structural features that distinguish different fibril polymorphs. Our analysis uncovered conserved and polymorph-specific features within alpha-synuclein fibrils. Notably, one site was conserved between *ex vivo* multiple system atrophy and *in vitro* H50Q mutant fibrils, suggesting the latter’s potential utility in drug discovery. Druggability predictions further prioritized ligandable sites across fibrils. Together, our findings establish a structure-based framework for identifying disease-relevant features and mapping them onto suitable *in vitro* models, guiding the rational development of polymorph-specific diagnostics and therapeutics for amyloid-related disorders with improved translational potential.

## Introduction

Neurodegenerative diseases (NDs) are characterized by the progressive loss of neurons, leading to impairments in memory, cognitive, and motor functions^1^. A defining hallmark of NDs is the misfolding and aggregation of soluble proteins into amyloid fibrils^2^, which deposit as extracellular plaques (e.g. Amyloid plaques)^3^ or in intracellular inclusions (e.g. Lewy bodies)^4^. Synucleinopathies and tauopathies are two sub-groups of NDs, marked by intercellular inclusions of aggregated alpha-synuclein (aSyn)^5^ and hyperphosphorylated tau^6^, respectively. Among NDs, Alzheimer’s disease (AD) and Parkinson’s disease (PD) are the most prevalent^7,8^, with the number of affected patients projected to reach 162 million worldwide by 2050^9^. Despite their growing societal impact, no preventive or disease-modifying therapies are currently available^10^.

Amyloid fibrils play a key role in the formation and spreading of ND’s pathology through several mechanisms. These include catalyzing monomer aggregation in a nucleation-dependent manner^11,12^, cell-to-cell propagation of pathology^13–17^ and inducing neurotoxicity by disrupting cellular membranes^18^, altering ion homeostasis and cellular proteostasis, and/or promoting oxidative stress^19–22^. Therefore, targeting amyloid fibrils offers opportunities to intervene with multiple pathogenic processes that contribute to disease development and progression^23–28^.

Advances in cryogenic electron microscopy (cryo-EM) have enabled determining the structure of amyloid fibril at near-atomic detail, revealing diverse architectures in both *in vitro* and *ex vivo* preparations^29–33^. These studies underscore the influence of aggregation conditions on fibril conformation^31,34^, demonstrating that a single protein can form fibrils of different structures (i.e., polymorphs)^35–37^–a phenomenon known as amyloid polymorphism^38^. Comparative cryo-EM structural analyses have revealed disease-associated fibril polymorphs across NDs^33^. In tauopathies, such as AD^39,40^, chronic traumatic encephalopathy (CTE)^41^, and Pick’s disease^42^, a unique tau fibril fold has been resolved for each condition (**Supplementary Fig. 1**A-D). Similarly, in synucleinopathies, including PD, dementia with Lewy bodies (DLB)^43^, multiple system atrophy (MSA)^37^, and juvenile-onset synucleinopathy (JOS)^44^, are each associated with structurally distinct aSyn fibrils (**Fig. 1A-D**). These polymorphs differ in protofilament number, fold and inter-protofilament interfaces, resulting in divergent surface topologies. Such structural features offer opportunities to develop polymorph-selective binders and disease-specific anti-amyloid therapeutics and diagnostics.

**Figure 1.**
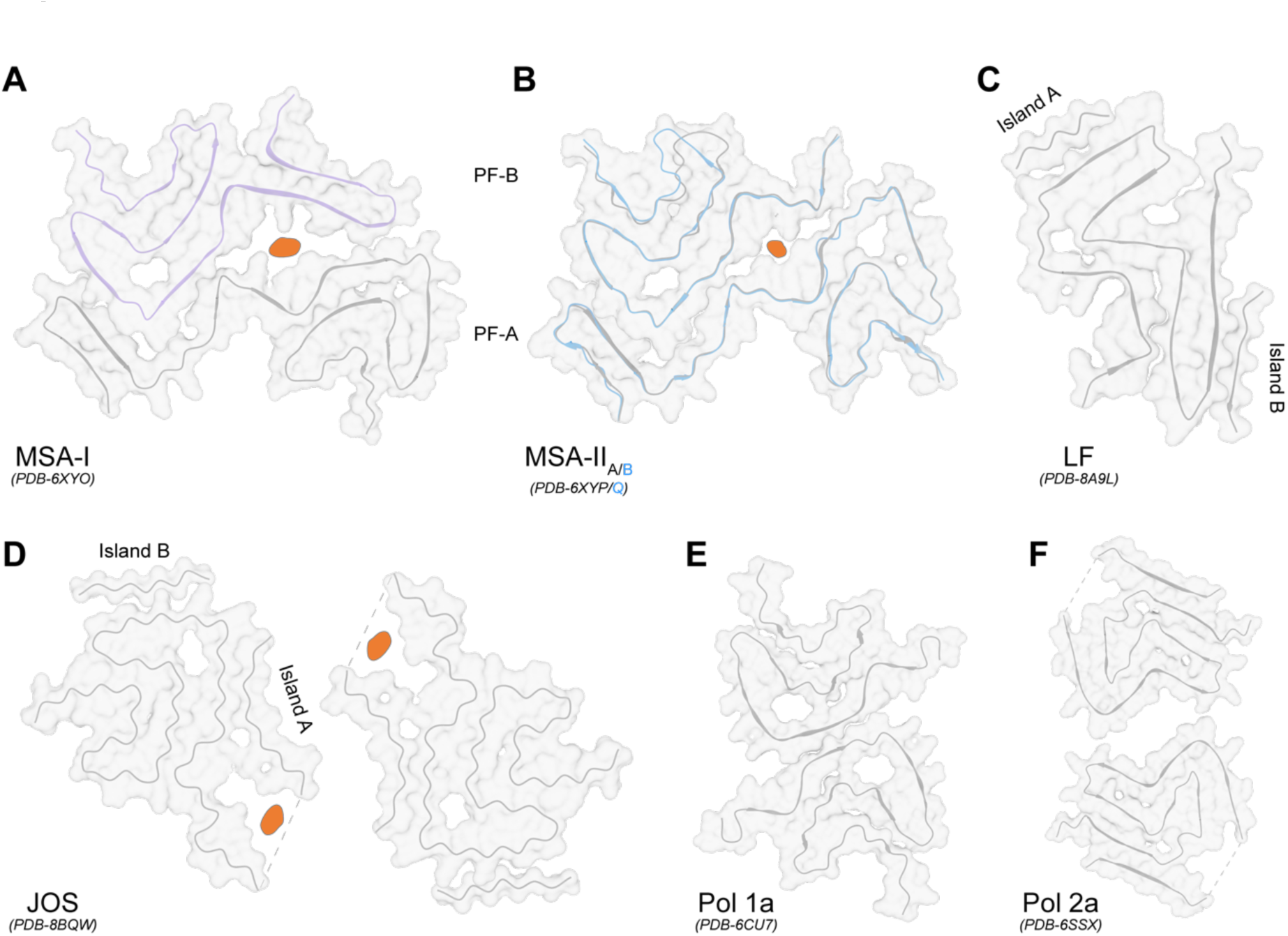
Representative alpha synuclein (aSyn) cryo-EM fibril structures. **(A-D)** Cryo-EM structures of ex vivo aSyn fibrils extracted from postmortem brain tissue of patients with multiple system atrophy^37^ (MSA; **A, B**), Parkinson’s disease (PD) and dementia with Lewy bodies^43^ (DLB; **C**) and juvenile-onset synucleinopathy^44^ (JOS; **D**). MSA fibrils consist of two asymmetric protofilaments, with protofilament A (PF-A) larger than protofilament B (PF-B). A non-proteinaceous density (orange) is present at the inter-protofilament interface and is shared between the MSA and JOS fibrils. **(E, F)** Cryo-EM structures of *in vitro* aSyn fibril polymorphs assembled from recombinant, human wild-type aSyn: polymorph 1a (Pol 1a; **E**) and polymorph 2a (Pol 2a; **F**). Both *in vitro* fibrils are composed of two symmetrical protofilaments. In Pol 1a, the protofilaments interact via a hydrophobic steric zipper^35^, whereas in Pol 2a the interaction is mediated by electrostatic salt-bridges^36^.

Although amyloid proteins readily aggregate *in vitro*, reproducing pathology-relevant fibril folds remains challenging. For example, recombinant wild-type (WT) aSyn fibrils adopt polymorphs that differ from those derived from synucleinopathy patients (**Fig 1**). Both *in vitro* WT and *ex vivo* MSA fibrils may comprise two protofilaments; however, the symmetric protofilaments in the *in vitro* fibrils interact via a hydrophobic steric zipper^35^ (**Fig 1E**) or electrostatic salt-bridges^36^ (**Fig. 1F**). In contrast, *ex vivo* MSA fibrils exhibit an extended interface between two asymmetric protofilaments, with a cavity that harbors an unidentified, non-proteinaceous density^37^ (**Fig. 1A, B**).

Additionally, *ex vivo* MSA fibrils, like other patient-derived fibrils, often contain unidentified co-factors and post-translational modifications (PTM), including acetylation, phosphorylation, and ubiquitination, which may influence fibril formation^37,45–47^. The absence of these biochemical features in *in vitro* aggregation conditions likely hinders the replication of pathological fibril polymorphs. Even when seeded amplification assays are employed, *ex vivo* fibrils give rise to structures resembling *in vitro* fibrils^48^. Notably, in the case of AD and CTE tau fibrils, their pathological fibril structure could be successfully generated only via the *de novo* aggregation of truncated tau proteins^49^ (**Supplementary Fig. 1**E, F).

Small molecules that bind amyloid fibrils hold promise for both diagnostics and therapeutics. Most discovery efforts have relied on high-throughput screening and optimization of know amyloidogenic dyes, such as Congo-red and thioflavin T (ThT)^50–52^. While several compounds have been identified, many lack target specificity. For instance, tracers developed for imaging amyloid plaques in AD patients, including BF-227^53^ and Pittsburgh compound B (PiB)^54^, also bind aSyn aggregates *in vitro*^55,56^ and in MSA patients^51,57^, likely due to shared structural motifs in their dye-based scaffolds^50,58^. More recently, an aSyn fibril-specific tracer (F0502B) was identified through an intensive screening and optimization process^59^, however, it failed to distinguish between different aSyn fibril polymorphs^60^. Despite these limitations, a few amyloid tracers have advanced to (pre)clinical diagnostic studies, although their selectivity remains a concern^51,56,61^.

Towards addressing the limitations outlined above, we developed a computational pipeline, FibrilSite, to identify shared and distinct surface features across selected fibril sites, based on geometric and chemical properties (**Fig. 2**). Focusing on sites from *ex vivo* aSyn fibrils, we assessed their similarity to those found in *in vitro* aSyn fibrils and in *ex vivo* fibrils formed by other amyloidogenic proteins. The conserved sites identified through this analysis can then be employed in computational drug-screening workflows, to prioritize candidate molecules for downstream experimental optimization^62,63^. To our knowledge, this represents the first comprehensive attempt to determine the extent to which *in vitro* generated fibrils could serve as reliable tools to discover or validate aSyn fibrils binders. This framework addresses key challenges in developing polymorph-and disease-specific diagnostics and therapeutics, including the rational selection of ligandable sites to achieve polymorph specificity and the strategic use of *in vitro* fibrils in drug development, particularly when *ex vivo* fibril folds cannot be reliably reproduced *in vitro*.

**Figure 2.**
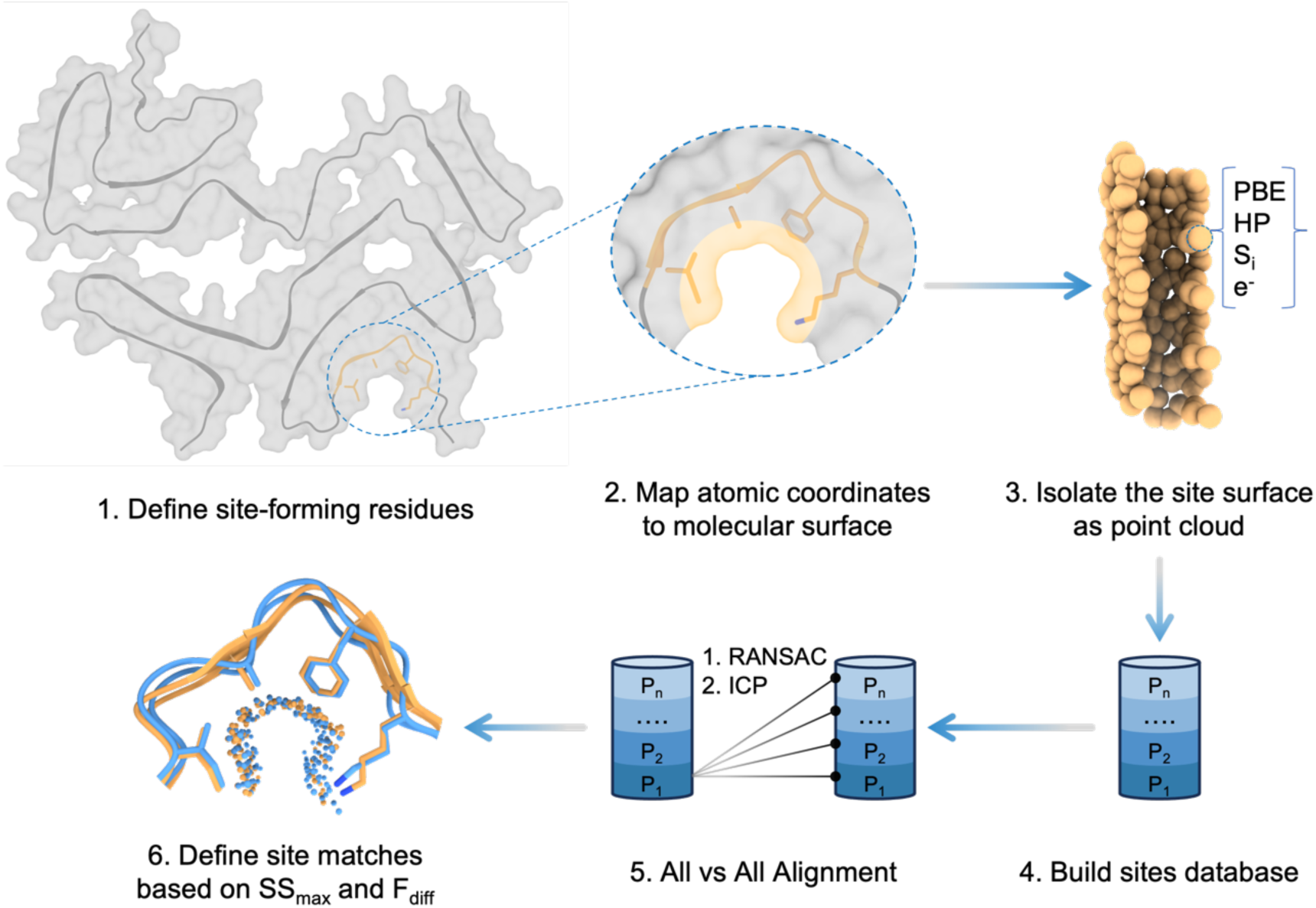
FibrilSite workflow for identifying conserved surface sites across structurally diverse amyloid fibrils. The FibrilSite workflow consists of six steps: (1) Site-forming residues are defined based on structural distinctiveness; (2) their atomic coordinates are mapped onto the fibril’s molecular surface; (3) each site is extracted as a point cloud and featurized with geometric and physicochemical surface properties, including Poisson–Boltzmann continuum electrostatics (PBE), hydrophobicity (HP), shape index (S_i_) and hydrogen bond donors/acceptor potential (e^-^) (Methods: FibrilSite, Defining fibril sites); (4) a database of 62 sites from 19 cryo-EM fibril structures is compiled; (5) all-vs-all alignments are performed using point cloud registration algorithms, including Random Sample Consensus (RANSAC) and Iterative Closest Point (ICP) (Methods: FibrilSite, Sites alignment); (6) similar sites are identified based on two metrics: surface overlap (SS_max_) and surface feature difference (F_diff_), which together assess the geometric and physicochemical similarity between aligned sites.

## Results

### Computational comparison of amyloid fibril site surfaces

To explore whether the distinct structural features of fibril sites can serve as markers for polymorph differentiation, we defined 62 sites from 19 fibril structures, including alpha synuclein (aSyn), tau, amyloid beta (Aβ), prion protein (PrP) and transmembrane protein 106B (TMEM106B). These sites were extracted and analyzed using our computational pipeline, FibrilSite (**Fig. 2**). Briefly, after defining the site-forming residues, each site was isolated as point cloud and annotated with geometric and physicochemical features, including electrostatics, hydrophobicity, hydrogen bond donors/acceptors, and shape index^64^.

To identify similar sites, we performed an all-vs-all alignment using point cloud registration algorithms, including Random Sample Consensus (RANSAC) and Iterative Closest Point (ICP), guided by local surface feature similarity. Site similarity was evaluated using two metrics: (i) SS_max_, which quantifies the maximum surface overlap between aligned sites while accounting for site size; and (ii) F_diff_, the Euclidean distance between corresponding surface features of aligned sites. Sites were considered similar if they exhibited both high SS_max_ and low F_diff_ values, indicating substantial surface overlap and minimal feature divergence.

### Identification of conserved and distinct surface sites in *ex vivo* amyloid fibrils

Given the promiscuity of amyloid-binding small molecules and the limited understanding of their binding sites, we systematically analyzed 27 defined sites in *ex vivo* fibrils: 15 from aSyn, 4 from tau, 2 from amyloid beta 42 (Aβ_42_), 2 from transmembrane protein 106B (TMEM106B), and 4 from prion protein (PrP), with a focus on identifying unique aSyn-specific sites (**Fig. 3, Supplementary Fig. 2**).

**Figure 3.**
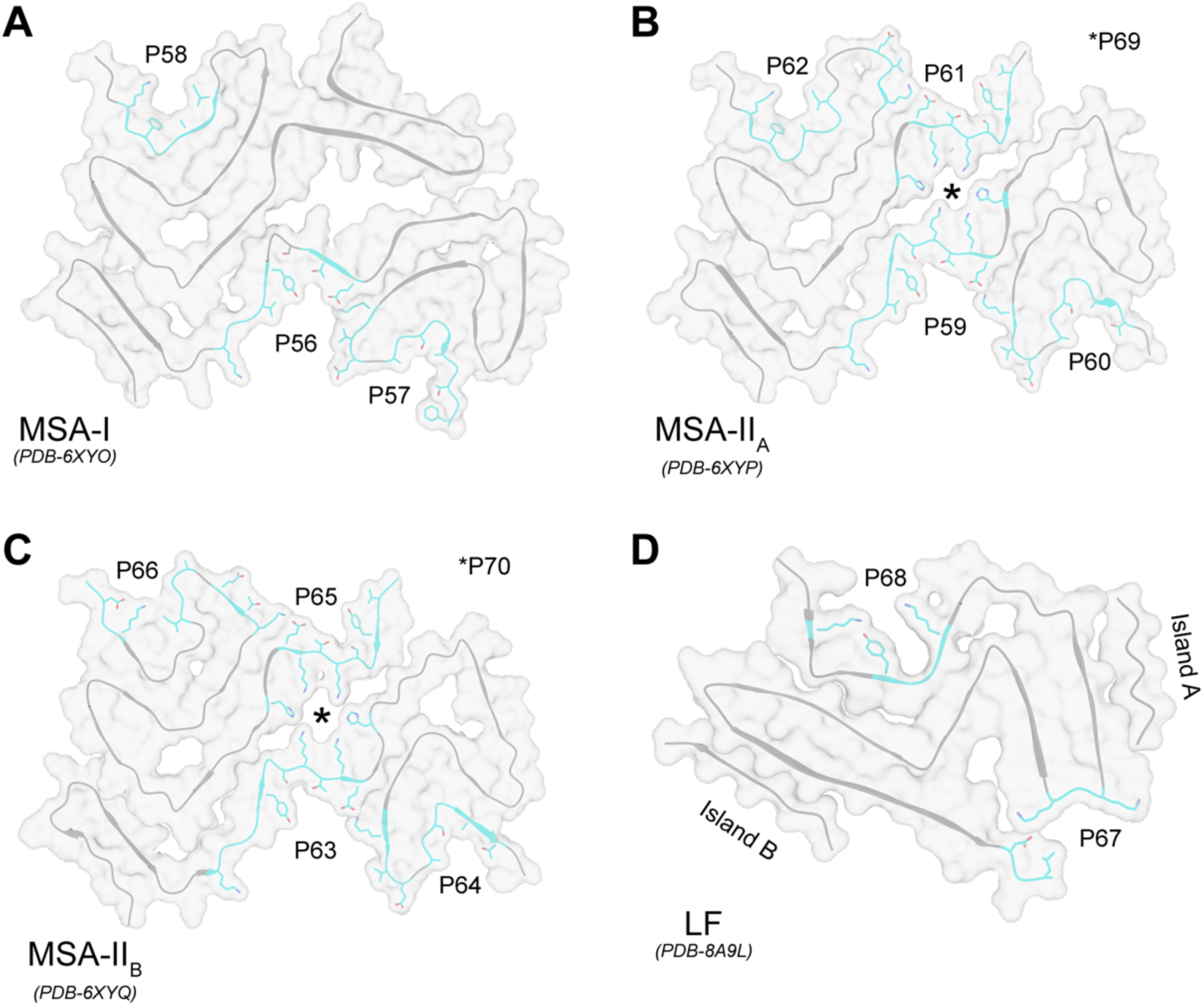
Defined sites from *ex vivo* alpha-synuclein (aSyn) fibrils. A total of fifteen sites were defined across four *ex vivo* aSyn fibril structures from patients with multiple system atrophy (MSA)^37^, Parkinson’s disease (PD) and dementia with Lewy bodies (DLB)^43^. Three MSA fibril polymorphs were analyzed: three sites were defined in MSA Type-I **(A)**, and five in each MSA Type-II variant^37^ **(B,C)**.The asterisk (*) indicates the central cavity site in Type-II fibrils, which is occupied by non-proteinaceous density. The Lewy fold (LF), associated with PD and DLB, contained two defined sites^43^ **(D)**. Site-forming residues are shown as cyan sticks on one fibril layer of each structure.

We first analyzed cryo-EM structures of *ex vivo* WT aSyn fibrils from patients with MSA, PD and DLB, to compare surface similarity between structurally divergent polymorphs with identical sequences. Two distinct folds were observed: a double protofilament fibril in MSA, featuring an N-terminus packed against a three-layered, L-shaped C-terminus; and a single-protofilament fibril–termed the Lewy Fold (LF)–in PD and DLB, characterized by a wave-like N-terminus with the C-terminus folded back onto it (**Fig. 1A-C**).

MSA fibrils consist of two asymmetric protofilaments, a larger protofilament A (PF-A) and a smaller protofilament B (PF-B), separated by a non-proteinaceous core. Structural variation within PF-B defined MSA fibril subtypes: in Type I fibrils, residues *Lys21-Gly36* were ordered and resolved; in Type-II fibrils, they were disordered. Further conformational differences in the *Ala78-Gln99* region subdivide Type-II fibrils into Type-II_A_ and Type-II_B_ (**Fig. 1A, B**).

In total, we defined 15 sites: three in MSA Type-I, five in MSA Type II_A_, five in MSA Type II_B_ and two in LF fibrils (**Fig. 3**). In MSA structures, two sites could be defined in PF-A across all fibril subtypes. The number of additional sites in PF-B varied by subtype, with one site in Type I, two in Type II_A_, and two in Type II_B_. Additionally, the central cavity site–occupied by a non-proteinaceous density–was included in both Type II subtypes.

Structural alignments confirmed the distinction between the two fibril folds, with similarities observed only among MSA fibril subtypes (**Fig. 4**). Two conserved sites with identical sequences were shared across MSA fibrils. The first, located in PF-A (*Ala85-Phe94*), exhibited greater surface overlap (SS_max_ = 0.89) and lower surface feature difference (F_diff_ = 0.31) between Type-II_A_ (P60) and Type-II_B_ (P64), compared to Type-I (P57), consistent with PF-A conservation within Type-II fibrils (**Fig. 4A**). The second site, located in PF-B (*Ile88–Lys96*) was conserved between Type-I (P58) and Type-II_A_ (P62) fibrils with SS_max_ = 0.87 and F_diff_ = 0.27 (**Fig. 4B**).

**Figure 4.**
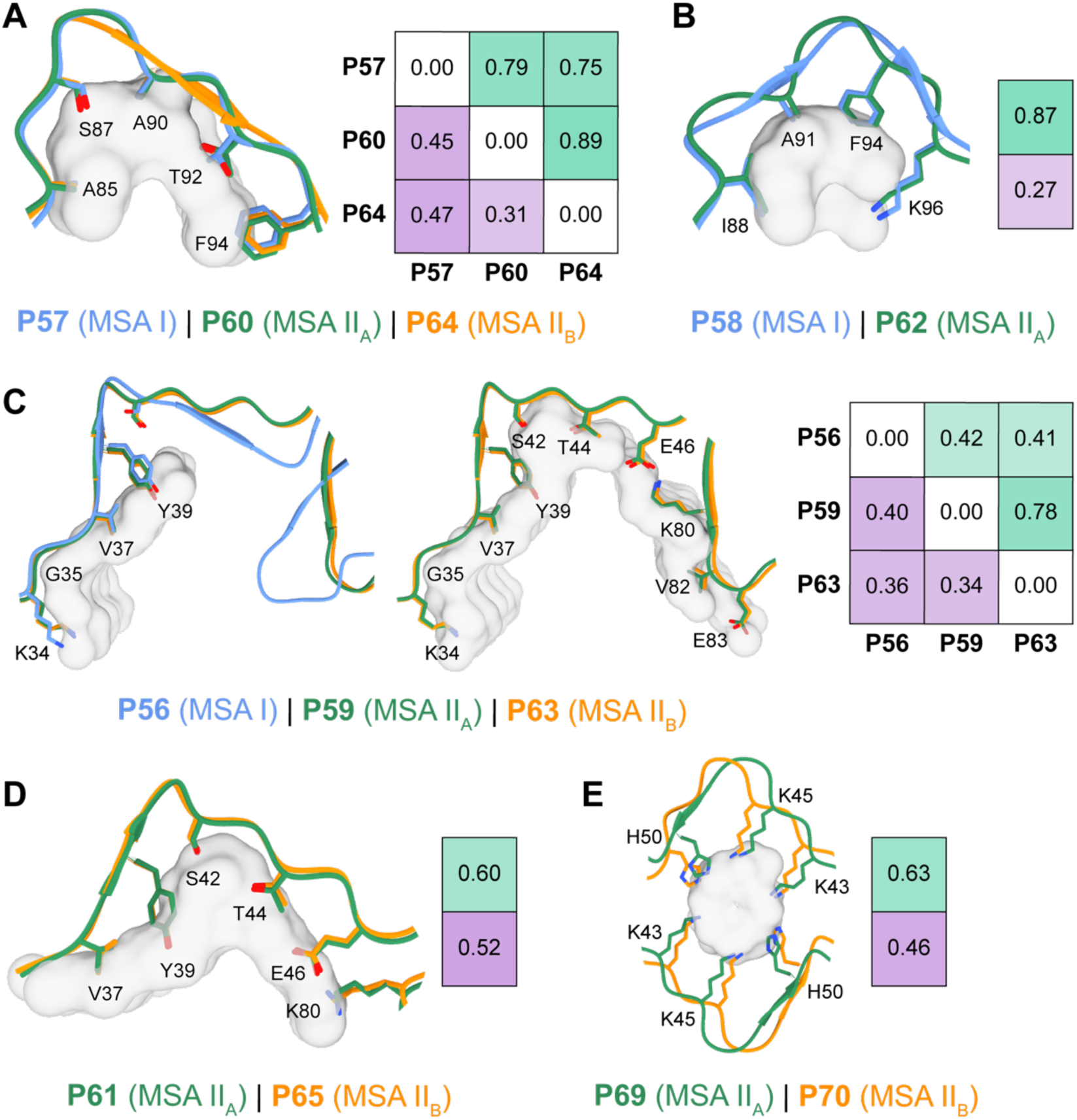
Conserved sites among multiple system atrophy fibril structures. Site matches were identified by aligning 15 sites from *ex vivo* alpha-synuclein (aSyn) fibril structures from patients with multiple system atrophy (MSA, 13 sites) and Parkinson’s disease (PD)/dementia with Lewy bodies (DLB; Lewy fold [LF], 2 sites), in an all-vs-all manner based on surface feature similarity. Features included electrostatics, hydrophobicity, hydrogen bond donors/acceptor potential, and shape index. For each matched pair, the site surface overlap (SS_max_, green shading, higher = better) and feature distance (F_diff_; Euclidean distance between aligned surface features; purple shading, lower = better) are shown. **(A)** A conserved site in protofilament A (PF-A) across all MSA fibrils, formed by residues *Ala85-Phe94*. **(B)** A shared site in PF-B of MSA Type-I and Type-II_A_ fibrils, formed by *Ile88-Lys96*. **(C)** A site in PF-A present in all MSA fibrils, defined by *Lys34-Glu46 and Lys80-Glu83*, showing extensive overlap in Type-II fibrils and partial overlap with Type-I fibrils (residues: *Lys34-Tyr39*). **(D)** A site in protofilament B (PF-B) of MSA Type-II fibrils, formed by *Val37-Glu46 and Lys80*. **(E)** A cavity site located in the core of MSA Type-II fibrils, occupied by non-proteinaceous density. Site-forming residues are shown as sticks; shared surface points are rendered in grey.

A third conserved site, located in PF-A (*Lys34-Glu46 and Lys80-Glu83*) was identified across all MSA fibrils. Although this site was highly conserved in Type-II fibrils (II_A_: P59, II_B:_ P63), its similarity with Type-I fibrils (P56) was limited to the *Lys34-Tyr39* region due to conformational differences in PF-A (**Fig. 4C**). Notably, the conformation of residues (*Val37-Glu46 and Lys80*) was conserved across multiple contexts in Type-II fibrils: between PF-A sites (P59, P63; **Fig. 4C**), between PF-B sites (P61, P65; **Fig. 4D**), and also between sites across PF-A and PF-B (**Supplementary Fig. 3**A).

Another conserved site corresponded to the Type-II fibrils central cavity, which harbors non-proteinaceous matter (**Fig. 4E**). All shared MSA sites had identical amino acid composition and substantial surface alignment, highlighting the internal structural diversity among MSA fibril types while reinforcing their uniqueness relative to LF fibrils.

The MSA sites conserved across all fibril subtypes represent potential targets for pan-MSA diagnostic and therapeutic strategies (**Fig. 4A, B**). Additionally, Type-II specific sites (**Fig. 4D, E**) may enable subtype-selective interventions, particularly for monitoring disease progression, as the Type-I to Type-II fibril ratio has been linked to disease duration^55^. Interestingly, some conserved MSA sites included residues such as Y39 and S87, which have been reported to undergo phosphorylation or nitration in disease^37,46,65,66^. However, a few studies suggested that the abundance of these modifications within fibrils is thought to be low^37,43^.

Sites with identical sequence and conformations helped refine the interpretation of site surface overlap (SS_max_) and surface feature difference (F_diff_) metrics, establishing a benchmark for site similarity: SS_max_ ≥ 0.5 and F_diff_ ≤ 0.6.

Next, we assessed surface similarity between sites from *ex vivo* aSyn fibrils and those of other amyloidogenic proteins (**Fig.2, Supplementary Fig. 2**). Due to differences in protein sequences and fibril polymorph conformation, no cross-protein site matches met our similarity criteria, highlighting the uniqueness of fibril surface features–not only among different amyloidogenic proteins, but also across polymorphs of the same protein (i.e., aSyn). Together, these findings support the potential of structurally defined fibril sites for polymorph- and disease-specific diagnostics and therapeutics.

### Comparative analysis of *in vitro* and *ex vivo* aSyn fibril surface sites

Next, we sought to identify sites shared between *ex vivo* and *in vitro* aSyn fibrils to assess the suitability of *in vitro* fibrils for screening pathologically relevant chemical matter. A major challenge in targeting pathological fibrils is the limited availability of patient-derived material, which is likely the most biologically relevant substrate for drug discovery and optimization. Reproducing pathology-relevant fibril folds *in vitro* has been largely unsuccessful, with the exception of *ex vivo* AD and CTE tau fibril folds^49^. An alternative strategy that can support drug development efforts involves computationally identifying specific features of brain-derived aggregates that are reproduced in one or more *in vitro*-generated fibrils, which could provide an intermediate solution for identifying disease-relevant fibril binders.

To explore this approach, we defined 35 sites from *in vitro* aSyn fibril structures assembled from recombinant human wild-type (WT) aSyn, disease-associated familial mutations and post-translationally modified variants (**Supplementary Fig. 4**). These sites were then compared to those previously defined in *ex vivo* aSyn fibrils (**Fig. 3**).

Among these, only two sites from the H50Q mutant fibril (P50 and P83) exhibited close structural similarity to a site (P66) in MSA Type-II_B_ fibrils. While minor backbone differences were observed, the side-chain arrangements were highly similar, yielding high surface overlap for P50 (SS_max_ = 0.69; **Fig. 5A**). All three sites share an identical sequence; however, the terminal residue *Asp98* is unresolved in P83, contributing to a shallower topology and reduced surface overlap with P66 (SS_max_ = 0.55; **Fig. 5A**). This match demonstrates that certain *in vitro* fibril preparations can replicate structural features present in brain-derived fibrils.

**Figure 5.**
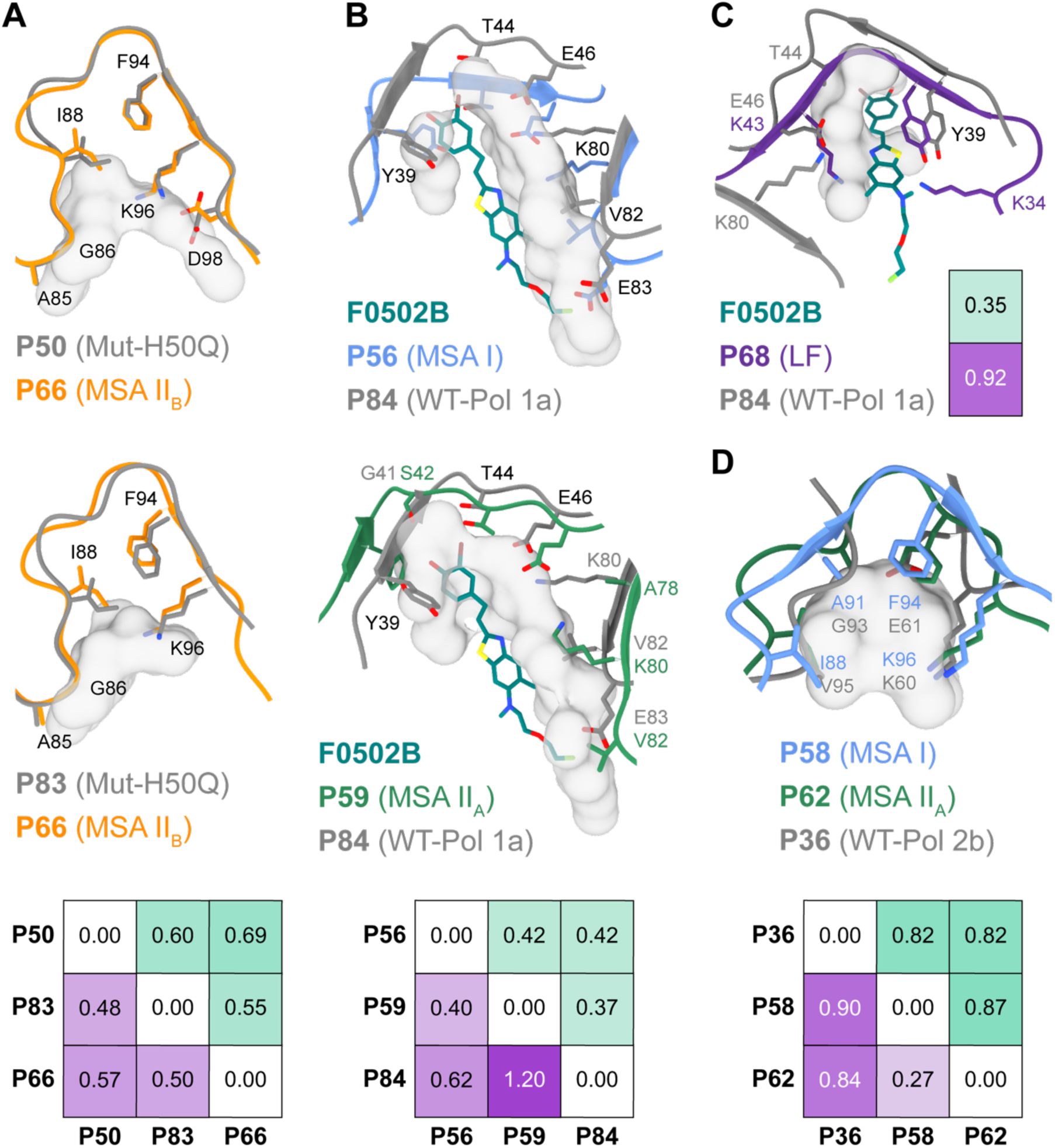
Shared sites between *ex vivo* and *in vitro* alpha synuclein (aSyn) fibrils. Site matches were identified by aligning 50 sites, including 15 from *ex vivo* (Fig. 3) and 35 from *in vitro* aSyn fibril structures (Supplementary Fig. 4). Alignments were performed in an all-vs-all manner based on surface feature similarity. Features included electrostatics, hydrophobicity, hydrogen bond donors/acceptor potential, and shape index. For each matched pair, the site surface overlap (SS_max_, green shading, higher = better) and feature distance (F_diff_; Euclidean distance between aligned surface features; purple shading, lower = better) are shown. **(A)** Alignment of a site in protofilament B (PF-B) of multiple system atrophy (MSA) Type-II_B_ (P66; formed by *Ala85-Asp98*) with two sites in the *in vitro* aSyn H50Q mutant fibrils (P50 and P83). **(B, C)** Alignment of the positron emission tomography tracer F0502B binding site (P84) in *in vitro* WT polymorph 1a fibrils with putative homologous sites in *ex vivo* MSA Type-I (P56), MSA Type-IIA (P59) **(B)** and Lewy Fold (LF; P68; **C**)**. (D)** Alignment of a site in PF-B of MSA Type-I (P58), MSA Type-II_A_ (P62) with a site in the *in vitro* WT Polymorph 2b fibrils (P36). Site-forming residues are shown as sticks; shared surface points are rendered in grey.

We next evaluated the alignments of the aSyn tracer F0502B binding site (P84; **Supplementary Fig. 4**D) from *in vitro* WT aSyn polymorph 1a fibrils with putative homologous regions in *ex vivo* aSyn MSA Type-I (P56), MSA Type-II_A_ (P59) (**Fig. 5B**) and the Lewy Fold (LF; P68) fibrils^59^ (**Fig. 5C**). Although F0502B labels pathological aSyn deposits in PD, DLB and MSA^59^, none of these alignments met our predefined site-similarity thresholds. Among MSA fibrils, P84 showed lower surface feature divergence with P56 in Type-I (F_diff_ = 0.62) than with P59 in in Type-II_A_ (F_diff_ = 1.2), with the latter showing substantial residue mismatches (**Fig. 5B**). Alignment with LF site (P68) resulted in the lowest surface overlap (SS_max_ = 0.35) and high surface feature divergence (F_diff_ = 0.92) (**Fig. 5C**). These findings, together with cryo-EM evidence of F0502B’s conformational adaptability across aSyn fibrils^60^, may explain its broad labelling across synucleinopathies. More broadly, they underscore the value of structural surface feature comparisons for identifying conserved fibril features.

Lastly, we evaluated the alignment with the highest surface overlap between *in vitro* WT and *ex vivo* aSyn fibrils. This involved P36 from *in vitro* WT polymorph 2b and two *ex vivo* MSA sites: P58 (Type-I) and P62 (Type-II_A_). Both alignments showed high surface overlap (SS_max_ > 0.8) but considerable surface features differences (F_diff_ = 0.90 and 0.82, respectively; **Fig. 5D**). While some residue aligned well–for example, *Lys60* in P36 aligned with *Lys96* in both MSA sites–others, such as *Glu61* in P36 aligning with *Phe94* in the MSA sites, introduced pronounced differences in charge and hydrophobicity, underscoring the divergence in surface chemistry despite geometric similarity.

Together, these comparisons highlight the structural differences between brain-derived and *in vitro*-prepared aSyn fibrils. Although shared sites were identified between MSA Type-II_B_ (P66) and the *in vitro* aSyn H50Q mutant fibril (P50 and P83), no high-confidence matches were found with *in vitro* WT aSyn fibrils commonly used in research. These findings illustrate the difficulty of reproducing disease-relevant structural features *in vitro* and support the use of computational strategies to identify conserved surface sites for structure-guided drug design. They further underscore the importance of accelerating efforts to reproduce pathology-associated fibril structures *in vitro* as essential tools to drive drug discovery and development of disease-specific imaging agents to track amyloid formation in the brain.

### Druggability assessment of fibril sites

To assess the druggability of the defined fibril sites, we employed P2Rank, a machine learning-based ligand-binding site predictor^67^. Given the atypical geometry and periodic topology of amyloid fibrils compared to globular proteins, we modified selected parameters in the default configuration to improve site detection sensitivity (see Methods, **Supplementary Table 1**). Additionally, because the defined sites often exceed the size typically occupied by a single ligand, P2Rank occasionally identified subregions within a given site as discrete pockets. Of the 62 sites defined across 19 cryo-EM fibril structures, P2Rank predicted 41 pockets across 38 sites, spanning 18 fibrils. No pockets were identified in the *in vitro* aSyn G51D mutant fibril. Identified pockets were subsequently evaluated using P2Rank’s predicted ligand-binding probability score (range: 0 (non-druggable) to 1 (druggable)). Applying a threshold of 0.7, 26 out of 41 pockets were classified as druggable (**Fig. 6**).

**Figure 6.**
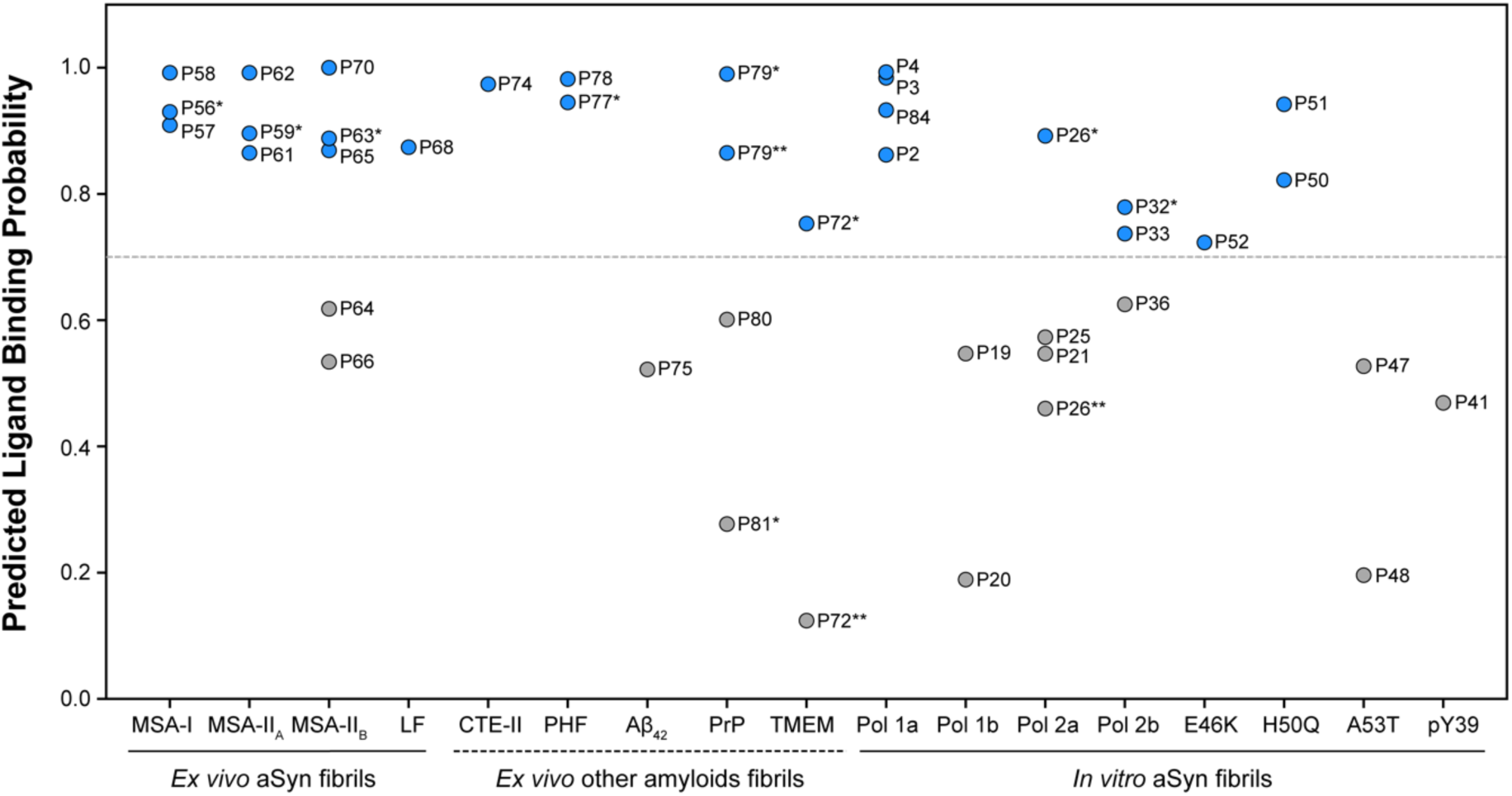
Druggability analysis of defined fibrils sites using P2Rank. Druggability of defined fibril sites was assessed using P2Rank^67^. Predicted ligand-binding probabilities (ranging from 0 = non-druggable to 1 = druggable) are shown for 41 identified pockets. To accommodate the atypical topological features of amyloid fibrils, the pocket definition algorithm was modified to improve detection of fibril sites. In some cases, P2Rank identified subregions within a single site, these are denoted by increasing numbers of asterisks (e.g., *, **) following the site name. A probability threshold of 0.7 (dashed line) was used to classify 26 pockets as druggable (blue) and 15 pockets as less-druggable (gray). Fibrils analyzed include *ex vivo* alpha-synuclein (aSyn) fibrils from MSA patients–Type I (MSA-I, PDB 6XYO), Type II_A_ (MSA-II_A_, PDB 6XYP), Type II_B_ (MSA-II_B_, PDB 6XYQ)–and from PD/DLB patients (LF, PDB-8A9L); *ex vivo* fibrils of other amyloidogenic proteins–Tau CTE-II (PDB 6NWQ), Tau PHF (PDB 7NRV), Aβ_42_ (PDB 7Q4M), PrP (PDB 7UMQ), TMEM106B (TMEM, PDB 7QVC); and *in vitro* aSyn fibrils, including wild-type polymorphs (Pol)–1a (PDB 6CU7, PDB 7WMM), 1b (PDB 6CU8), 2a (PDB 6SSX), 2b (PDB 6SST); familial mutants–E46K (PDB 6UFR), H50Q (PDB 6PES), A53T (PDB 6LRQ), and phosphorylated aSyn at *Tyr39*–pY39 (PDB 6L1U).

Focusing on *ex vivo* fibrils, all structures contained at least one druggable site, with the exception of Aβ_42_ fibrils. Notably, P78 site in tau paired helical filaments (PHF) was predicted to be druggable, consistent with cryo-EM studies identifying it as a binding site for EGCG^68^. Among *ex vivo* aSyn fibrils, one of the two defined sites in Lewy fold (LF) fibrils was detected and predicted to be druggable. In contrast, 11 out of 13 defined sites in MSA fibrils were identified by P2Rank, of which 9 were classified as druggable. Among these, the matched sites in Type I (P58) and Type-II_A_ (P62) fibrils (**Fig. 4B**), as well as sites in Type-II_A_ (P59 and P61) and Type-II_B_ (P63 and P65) fibrils (**Fig. 4D, Supplementary Figure 3**), exhibited similar druggability scores, reinforcing the expectation that structurally and chemically similar sites possess comparable ligand-binding potential.

Among *in vitro* aSyn fibrils, all but WT polymorph 1b and A53T mutant fibrils contained druggable pockets. In WT polymorph 1a, druggable sites included P84, which corresponds to the tracer F0502B binding pocket^59^, as well as P2 and P4, both of which were recently shown by cryo-EM studies to accommodate various small molecules^56,60^. These predictions align with experimental findings, providing confidence in the method’s potential for analyzing amyloid fibrils.

Despite P2Rank’s overall effectiveness in identifying druggable sites in amyloid fibrils, several discrepancies highlighted the limitations of current pocket prediction tools when applied to non-canonical structures such as amyloid fibrils. For example, among the matched sites in MSA Type I (P57), Type-II_A_ (P60), and Type-II_B_ (P64) fibrils (**Fig. 4A**), only P57 was predicted to be druggable, while P64 scored much lower and P60 was not detected. Similarly, for the matched sites between MSA Type-II_B_ (P66) and the *in vitro* aSyn H50Q mutant (P50) fibrils (**Fig. 5A**), P50 was predicted to be druggable, while P66 scored below the set threshold. Since these matched sites share identical sequences and similar surface properties, their divergent predictions highlight the need for pocket detection algorithms tailored to the periodicity and surface characteristics of amyloid fibrils.

Lastly, although P83 from the *in vitro* aSyn H50Q mutant fibrils also aligned with P66 from MSA Type-II_B_ (**Fig. 5A**), it was not identified as a pocket, likely due to its shallower topology. Taken together, P50 emerged as the only site from an *in vitro* aSyn fibril with both high predicted druggability and strong structural similarity to an *ex vivo* fibril site, underscoring its potential translational relevance. These findings highlight the structural diversity and druggability of many fibril sites and support the feasibility of developing disease-specific molecules based on fibril fold differentiation.

## Discussion

A major challenges in the rational design of amyloid-binding molecules lies in the limited availability of patient-derived fibrils and the structural divergence between *in vitro* and pathological fibril folds^33^. High-throughput drug screening often relies on *in vitro* fibrils, which frequently fail to replicate structural features of pathological conformations, leading to molecules with limited specificity^57,61,69^. Nevertheless, the discovery of F0502B – an aSyn fibril-specific tracer – using *in vitro* WT aSyn fibrils, demonstrates the potential of fibril-specific drug development through comprehensive screening and optimization^59^. Cryo-EM studies have revealed that F0502B induces a conformational rearrangement in the *in vitro* fibrils, creating a binding cavity resembling those found in *ex vivo* aSyn fibrils^59^. Also, its binding adaptability to different site conformations across aSyn fibrils^60^, likely accounts for its ability to label both *in vitro* and pathological assemblies. These findings underscore the need for strategies that bridge the gap between *in vitro* systems and disease-relevant fibril conformations.

To address this gap, we developed FibrilSite, a computational pipeline for comparing amyloid fibril surface sites based on geometric and physicochemical features (**Fig. 2**). Using this framework, we analyzed 62 sites from 19 fibril structures spanning multiple amyloidogenic proteins. Site alignments were performed using point-cloud registration algorithms guided by encoded surface properties, enabling quantification of surface overlap (SS_max_) and feature divergence (F_diff_). The pipeline is modular and expandable, supporting integration of new structures and binding sites.

This approach revealed the structural uniqueness of *ex vivo* aSyn fibril: no sites were shared between fibrils from MSA and PD/DLB patients, or with other amyloidogenic protein fibrils. Within MSA fibrils, we identified conserved sites shared across all polymorphs **(Fig 4A)**, suggesting potential pan-MSA targets. We also identified sites unique to Type-II fibrils **(Fig 4E)**, which could enable subtype-selective targeting. These distinctions may hold clinical relevance, as the Type-I/Type-II ratio is thought to correlate with disease duration^37^.

We further identified structurally conserved sites between *ex vivo* MSA Type-II_B_ fibrils (P66) and the *in vitro* aSyn H50Q mutant fibrils (P50 and P83; **Fig. 5A**), suggesting that certain pathological features can be recapitulated *in vitro*. These findings underscore the value of identifying shared fibril features between *ex vivo* and *in vitro* fibrils to guide ligand discovery. Computational and experimental screening efforts focused on these common features may improve translational relevance, particularly when pathological folds cannot yet be reproduced *in vitro*.

To prioritize fibril sites for ligand design, we evaluated their druggability using P2Rank^67^, a machine learning-based ligand-binding site predictor. We optimized selected parameters to account for the non-globular, periodic geometry of fibrils. Of 62 sites, 41 pockets were identified, 26 of which exceeded a ligand-binding probability of 0.7 and were considered druggable (**Fig. 6**). Structurally similar site pairs exhibited similar druggability scores, supporting the notion that chemically and geometrically similar surfaces tend to exhibit comparable ligand-binding potential. However, discrepancies were also observed, for example some structurally similar sites with identical sequences yielded differing predictions, underscoring the limitations of current pocket prediction tools when applied to amyloid fibrils and highlight the need for algorithms tailored to amyloid geometries.

Beyond surface features, other factors influence druggability include site accessibility and presence of PTMs. Fibril cryo-EM structures resolve only the ordered core, while disordered regions form a dynamic “fuzzy coat” surrounding the core, which may hinder ligand access^33,70–73^. Furthermore, pathological aggregates often contain PTMs (e.g., phosphorylation and nitration)^4,37,66^, which can alter the chemical environment of binding sites, potentially influencing ligand interactions and pathogenicity^74,75^.

Furthermore, Ligand binding modality itself further modulates selectivity and affinity. Cryo-EM studies of ligand-bound fibrils^56,59,60,68^, have revealed that specific ligands, bind diagonally across fibril layers, enabling inter-ligand π–π stabilizing interactions, whereas, less specific ligands, bind horizontally and lack such interactions^56^. These observations emphasize the complexity of ligand-fibril interactions and highlight key structural considerations for designing selective fibril-binding molecules.

In summary, our findings demonstrate the utility of computational approaches for identifying structurally conserved and druggable sites in pathological fibrils and mapping them onto *in vitro* models. This strategy enables the prioritization of targetable sites and the rational selection of appropriate *in vitro* systems to improve the translational potential of drug development efforts, especially in cases where disease-relevant fibril folds remain experimentally inaccessible. Cross-protein site comparisons facilitate polymorph- and disease-specific targeting while minimizing off-target effects. Combining advances in machine learning drug design against amyloids^76^ with structure-based directed targeting and high-resolution structural biology, the precision and effectiveness of fibril-targeting drug design can be substantially enhanced, paving the way for more selective diagnostics and therapeutics for neurodegenerative diseases.

## Data availability

All data for this work are freely accessible on Zenodo at https://doi.org/10.5281/zenodo.15192320

## Code availability

The code and scripts for FibrilSite, used in site definition, alignment, and analysis, are available online at https://github.com/A-Sadek/FibrilSite

## Acknowledgment

We thank SCITAS at EPFL for support in running our pipeline and analysis. We thank Evgenia Elizarova and Arne Schneuing for assistance with computational method development (Laboratory of Protein Design and Immunoengineering, Institute of Bioengineering, EPFL, Switzerland). We thank Lucas Burget assistance in running fibril site alignment on the high-performance computing cluster and Danaé Terrien-Ferey for assistance with identifying the fibril sites in amyloid fibrils from other amyloid proteins (Laboratory of Molecular and Chemical Biology of Neurodegeneration, École Polytechnique Fédérale de Lausanne (EPFL), Lausanne, Switzerland).

## Author information

Authors and Affiliations

**Laboratory of Molecular and Chemical Biology of Neurodegeneration, École Polytechnique Fédérale de Lausanne, Lausanne, Switzerland**

Ahmed Sadek, Hilal A. Lashuel

**Laboratory of Protein Design and Immunoengineering, Institute of Bioengineering, Ecole polytechnique fédérale de Lausanne, Lausanne, Switzerland**

Ahmed Sadek, Bruno E. Correia

**Weill Cornell Medicine Qatar, Education City, Qatar Foundation, Doha, Qatar**

Hilal A. Lashuel

**Department of Neurology, Weill Cornell Medicine, New York, NY, USA**

Hilal A. Lashuel

## Contributions

H.A.L., B.E.C., and A.S. conceived the project. A.S. developed FibrilSite algorithm and organized the GitHub repository. A.S. carried out the analysis and created the figures with input from all authors. All authors contributed to writing and revising the manuscript.

## Corresponding authors

Correspondence to Bruno E. Correia and Hilal A. Lashuel

## Ethics declarations

Competing interests

H.A.L is the founder and CEO of ND BioSciences, a spinoff from the Lashuel lab focusing on developing novel therapies and diagnostics for neurodegenerative diseases.

## Methods

### Computing molecular surfaces and surface features

Molecular surfaces and surface features were computed as described in [64]. Briefly, fibril-deposited PDB structures were protonated using Reduce^77^, triangulated with MSMS^78^ at a density of 3.0 and a water probe radius of 1.5 Å, then downsampled and regularized to a 1Å resolution using PyMesh^79^. Each vertex was assigned four features: Poisson–Boltzmann electrostatics, hydropathy, shape index, and hydrogen bond potential. Electrostatic properties were computed using APBS^80^, with charge values constrained to ±30 and normalized to ±1. Hydropathy values, based on the Kyte and Doolittle scale^81^, were assigned according to the amino acid identity of the vertex’s closest atom, ranging from −4.5 (hydrophilic) to +4.5 (hydrophobic) and normalized to ±1. The shape index quantified local curvature^82^ with values ranging from −1 (highly concave) to +1 (highly convex). Hydrogen bond donor and acceptor potentials were determined using a Gaussian-weighted hydrogen bond potential^83^, with values from −1 (optimal acceptor) to +1 (optimal donor) based on atomic proximity and orientation.

### FibrilSite, Defining fibril sites

Defined sites were manually extracted using a tailored algorithm incorporating point cloud and mesh handling packages, including PyMesh^79^, Open3D^84^, and Biopython^85^. The fibril atomic structure (PDB file) and its corresponding molecular surface (Ply file) were first loaded, and the fibril’s main axis was computed using eigenvector decomposition. Surface points were classified based on their orientation relative to this axis using the dot product, enabling the isolation of the lateral fibril surface. Site-forming residues served as anchor points, and their nearest surface vertices were identified to define the core site region. This region was subsequently expanded by including neighboring surface points within 5Å, retaining only those with similarly oriented surface normals (dot product > 0). Finally, a rim residue was manually designated, and excess peripheral points were trimmed to ensure accurate delineation of each site surface.

### FibrilSite, Sites alignment

Isolated site surfaces, represented as point clouds, were aligned in an all vs all manner using Random Sample Consensus (RANSAC) and Iterative Closest Point (ICP) algorithms implemented in Open3D^84^. First, the source and target sites were centralized in space by subtracting the mean coordinate value from each surface point for both sites. RANSAC then performed 20,000 iterations to align the source site to the target, prioritizing points with minimal surface feature differences and selecting the transformation with the most points within 1Å. Finally, ICP refined the alignment for optimal accuracy.

### Site Surface overlap metric (SS_max_)

SS_max_ quantifies the maximum overlap between two aligned site surfaces, providing a measure of structural similarity. It is defined as:

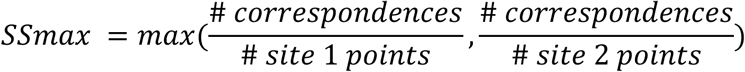

where correspondences refer to the number of matched points between the two surfaces after alignment. The metric ensures that the overlap is evaluated relative to both site surfaces, accounting for differences in size. A higher SS_max_ value indicates a greater degree of shared surface between the two sites.

### Assessment of fibril sites druggability with P2Rank

To assess the druggability of defined fibril sites, we used P2Rank (v2.5), a machine learning-based ligand-binding site predictor^67^. Because amyloid fibrils exhibit non-globular, periodic and structurally distinct topologies that diverge from the training data used to develop such models, we modified selected parameters in the default configuration to improve the algorithm’s sensitivity to fibril-specific features. These adjustments, summarized in **Supplementary Table 1**, enhanced the tool’s ability to capture the diversity of fibril surface environments and more effectively identify ligandable sites. Predicted pockets were evaluated based on P2Rank’s ligand-binding probability score (range: 0 = non-druggable, 1 = druggable), and classified as druggable if the score exceeded 0.7.

---

**Supplementary Table 1.**
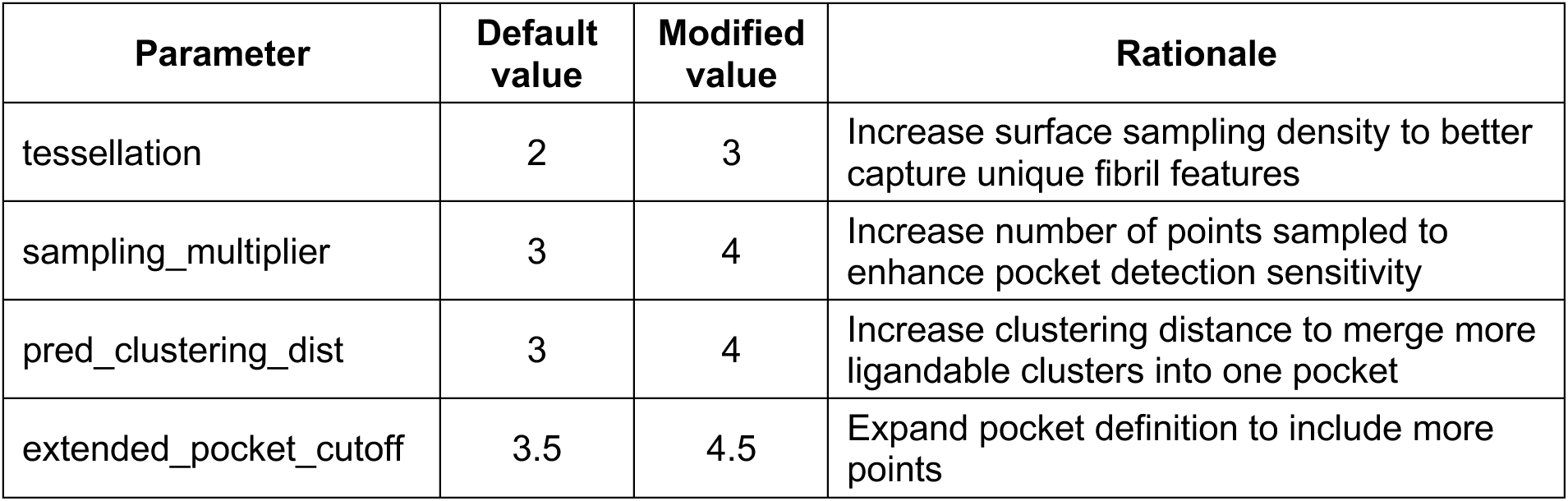
Modified P2Rank parameters for amyloid fibril pocket detection. Selected parameters in the default P2Rank^67^ (v2.5) configuration were adjusted to account for the distinctive geometry of amyloid fibrils. Modifications were made to increase surface sampling density and improve sensitivity to extended or shallow surface grooves characteristic of fibril fold architectures. The full configuration file is provided in the accompanying data repository.

## Supplementary figures

**Supplementary Figure 1.**
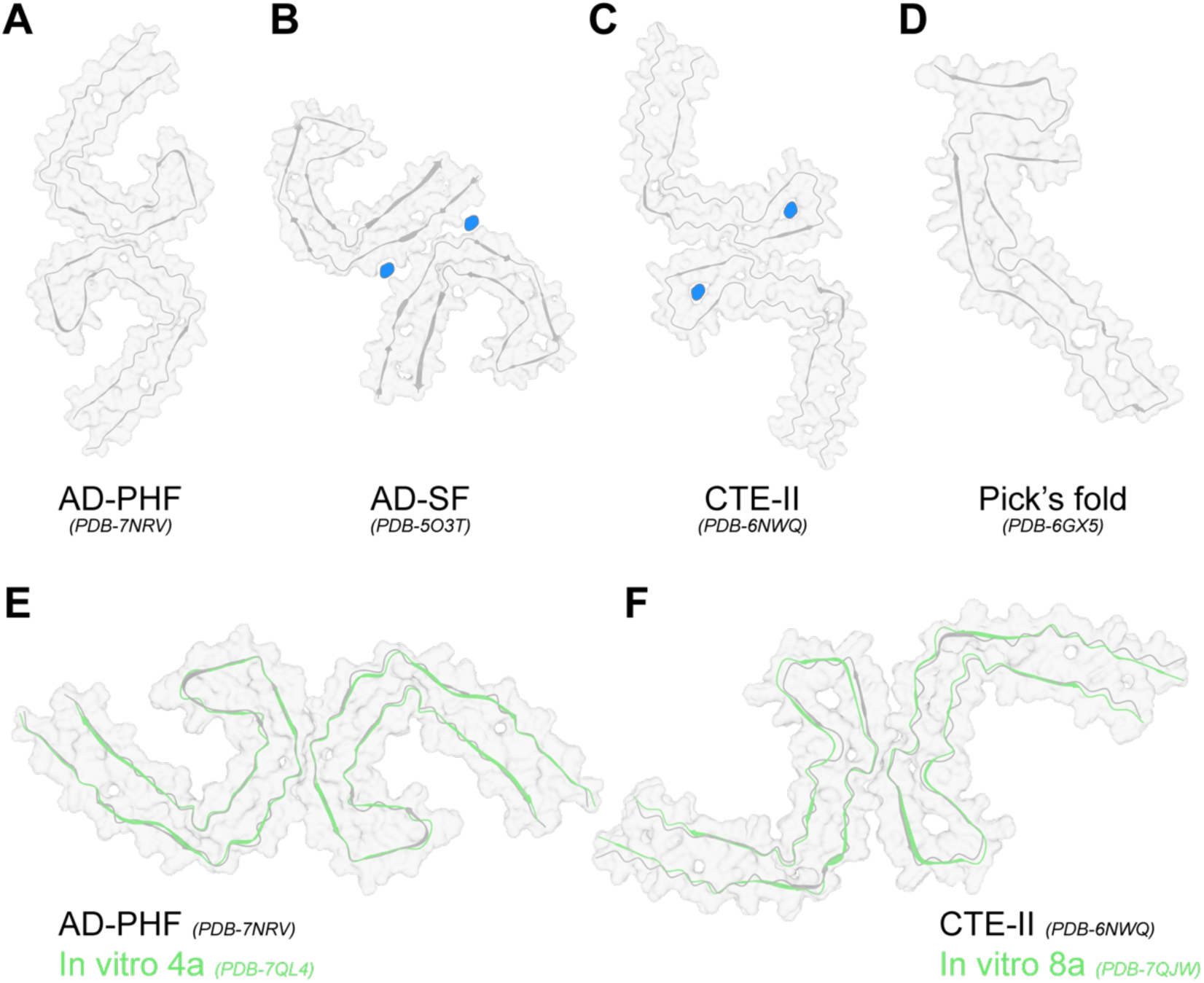
Representative tau fibril cryo-EM structures. **(A, B)** *Ex vivo* tau paired helical filaments (PHF) and straight filaments (SF) from Alzheimer’s disease (AD) patient brains^39,40^. **(C)** *Ex vivo* tau fibrils from chronic traumatic encephalopathy (CTE) patients^41^; two structural variants were identified, both composed of identical protofilaments with distinct inter-protofilament interfaces–Type-II fibrils are shown. **(D)** *Ex vivo* tau fibrils from Pick’s disease patients^42^. Blue densities correspond to potential cofactors, influencing the SF fold in AD and becoming entrapped in the CTE fibril protofilaments. **(E, F)** Overlay of *in vitro* generated fibrils^49^ (green) with their corresponding *ex vivo* counterparts (gray): AD-PHF fold **(E)** and CTE-II fold **(F)**.

**Supplementary Figure 2.**
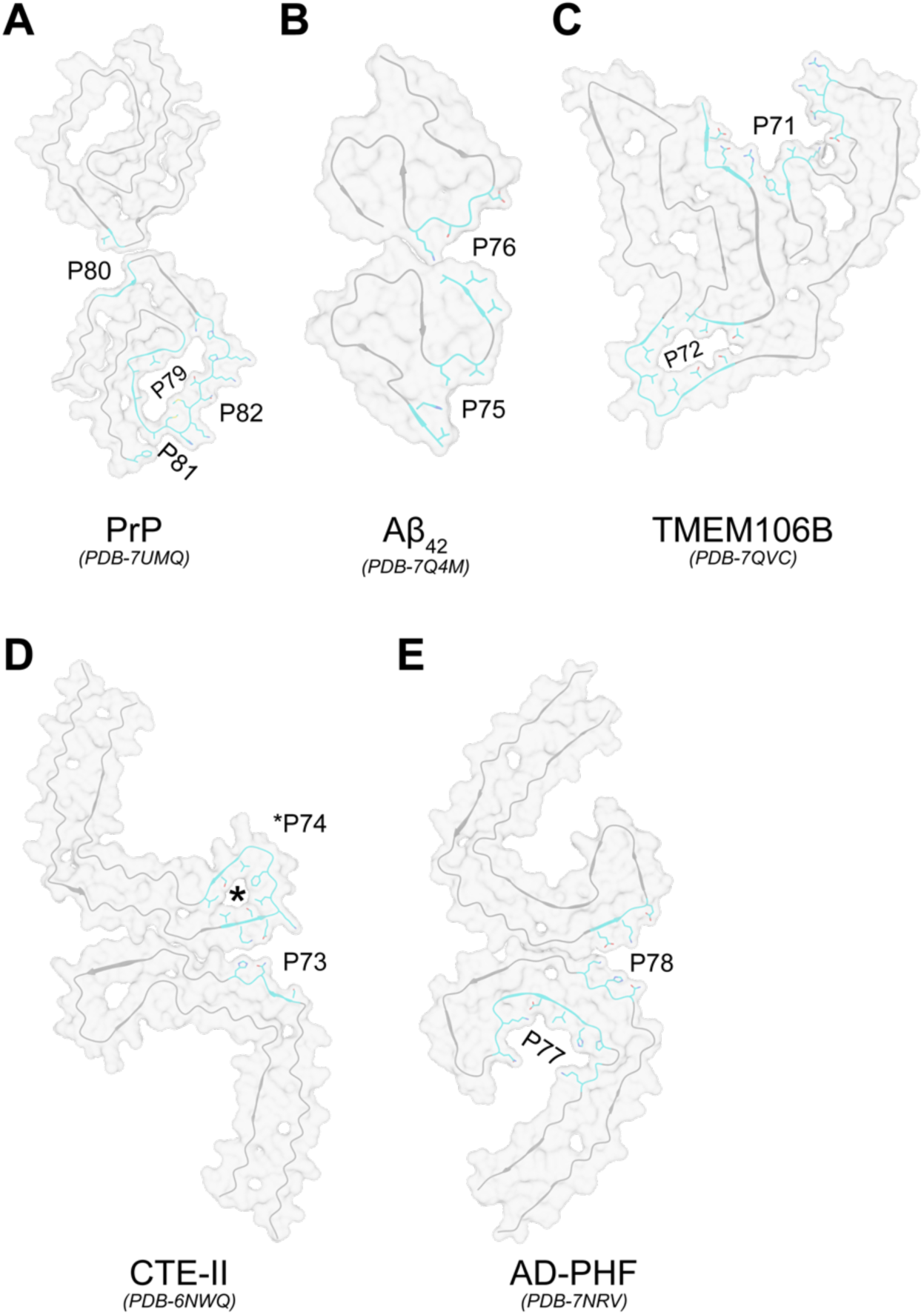
Defined sites from *ex vivo* amyloid fibrils of diverse origins. A total of twelve sites were defined across five *ex vivo* fibril structures representing different amyloidogenic proteins.**(A)** Four sites in prion protein (PrP) fibrils isolated from Gerstmann-Straussler-Scheinker disease patients^86^. **(B)** Two sites in in Type-II amyloid beta 42 (Aβ_42_) fibrils isolated from familial Alzheimer’s disease (AD) patients^87^. **(C)** Two sites in transmembrane protein 106B (TMEM106B) fold I fibrils isolated from AD patients^88^. **(D)** Two sites in type-II tau fibrils isolated from chronic traumatic encephalopathy (CTE) patients^41^. **(E)** Two sites in tau paired helical filaments (PHF) isolated from AD patients^40^. Site-forming residues are shown as cyan sticks on one fibril layer of each structure.

**Supplementary Figure 3.**
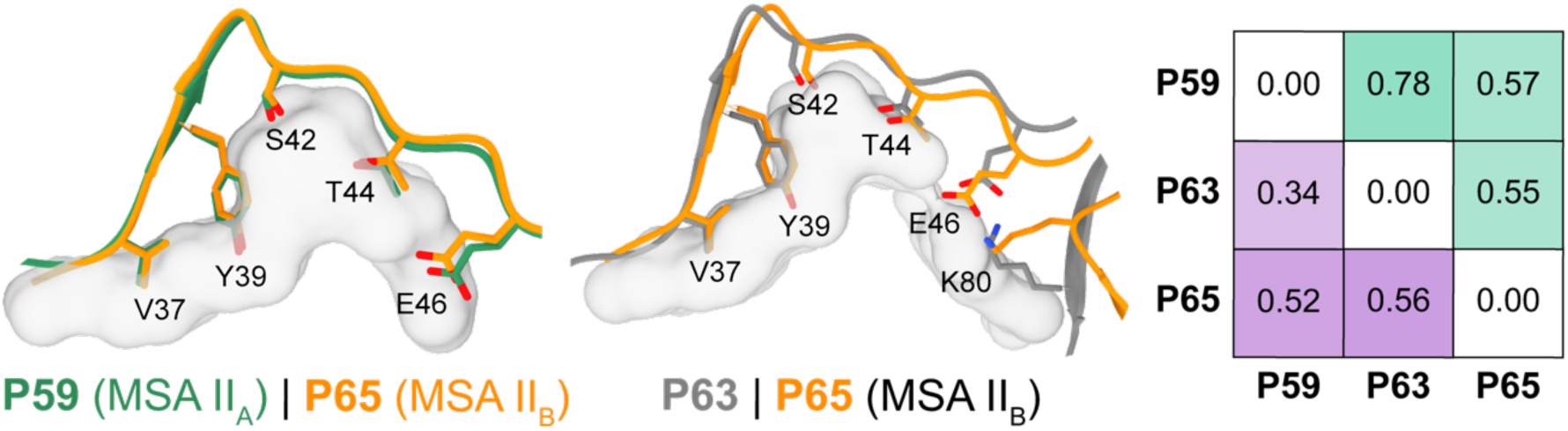
Conserved sites among *ex vivo* alpha-synuclein (aSyn) MSA fibrils. Site matches were identified by aligning 15 sites from *ex vivo* aSyn fibrils (Fig. 3) against one another. Alignments were performed based on surface feature similarity, including electrostatics, hydrophobicity, hydrogen bond donors/acceptor potential, and shape index. For each matched pair, the site surface overlap (SS_max_, green shading, higher = better) and feature distance (F_diff_; Euclidean distance between aligned surface features; purple shading, lower = better) are shown. A conserved site formed by *Val37-Glu46 and Lys80* in multiple system atrophy (MSA) Type-II fibrils, shared between protofilament A (PF-A) of Type-II_A_ (P59) and protofilament B (PF-B) of Type-II_B_ (P65), as well as between PF-A (P63) and PF-B (P65) of Type-II_B_. Site-forming residues are shown as sticks; shared surface points are rendered in grey.

**Supplementary Figure 4.**
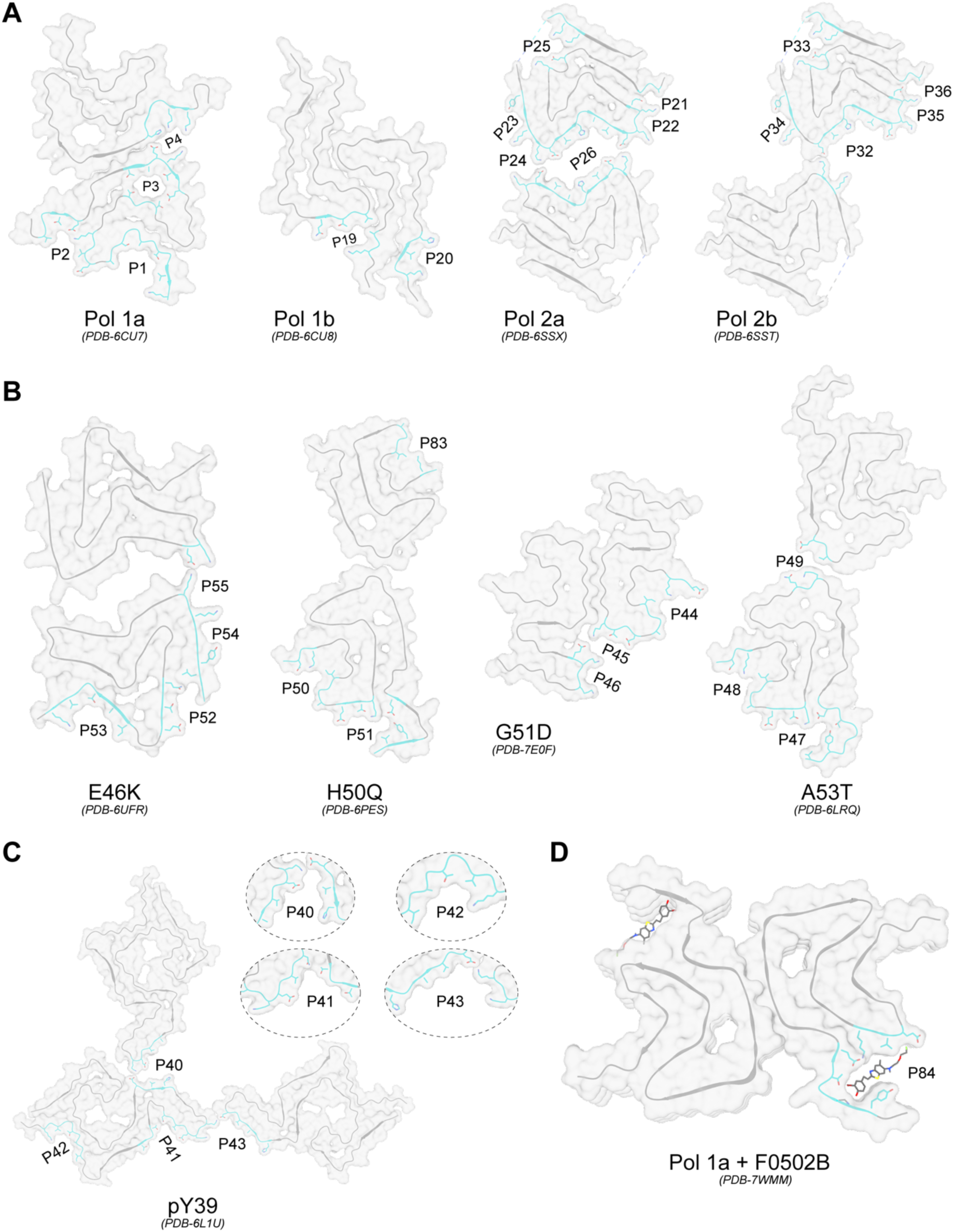
Defined sites from *in vitro* alpha-synuclein (aSyn) fibrils. A total of 35 sites were defined across ten *in vitro* aSyn fibril structures. **(A)** Seventeen sites were defined in four wild-type (WT) aSyn fibril polymorphs^35,36^ (Pol): four in Pol 1a, two in Pol 1b, six in Pol 2a, and five in Pol 2b. **(B)** Thirteen sites were defined in fibrils formed by aSyn variants harboring familial mutations (Mut): four in E46K fibril^89^, three in H50Q^90^, three in G51D fibril^91^, and three in A53T fibril^92^. **(C)** Four sites were defined in fibrils formed by aSyn phosphorylated at *Tyr39*^45^. **(D)** PET tracer F0502B binding site in WT Pol 1a. Site-forming residues are shown as cyan sticks on one fibril layer of each structure.

## References

1. Wilson, D. M. et al. Hallmarks of neurodegenerative diseases. Cell 186, 693–714 (2023).

2. Iadanza, M. G., Jackson, M. P., Hewitt, E. W., Ranson, N. A. & Radford, S. E. A new era for understanding amyloid structures and disease. Nat. Rev. Mol. Cell Biol. 19, 755–773 (2018).

3. Fiala, J. C. Mechanisms of amyloid plaque pathogenesis. Acta Neuropathol. (Berl.) 114, 551–571 (2007).

4. Fares, M. B., Jagannath, S. & Lashuel, H. A. Reverse engineering Lewy bodies: how far have we come and how far can we go? Nat. Rev. Neurosci. 22, 111–131 (2021).

5. Visanji, N. P., Lang, A. E. & Kovacs, G. G. Beyond the synucleinopathies: alpha synuclein as a driving force in neurodegenerative comorbidities. Transl. Neurodegener. 8, 28 (2019).

6. Zhang, Y., Wu, K. M., Yang, L., Dong, Q. & Yu, J. T. Tauopathies: new perspectives and challenges. Mol. Neurodegener. 2022 171 17, 1–29 (2022).

7. Andrade-Guerrero, J. et al. Alzheimer’s Disease: An Updated Overview of Its Genetics. Int. J. Mol. Sci. 24, 3754 (2023).

8. Zhu, J. et al. Temporal trends in the prevalence of Parkinson’s disease from 1980 to 2023: a systematic review and meta-analysis. Lancet Healthy Longev. 5, e464– e479 (2024).

9. Chopade, P., et al. Alzheimer’s and Parkinson’s disease therapies in the clinic. Bioeng. Transl. Med. 8, e10367 (2023).

10. Mortada, I. et al. Immunotherapies for Neurodegenerative Diseases. Front. Neurol. 12, 654739 (2021).

11. Nonaka, T., Watanabe, S. T., Iwatsubo, T. & Hasegawa, M. Seeded aggregation and toxicity of α-synuclein and tau: Cellular models of neurodegenerative diseases. J. Biol. Chem. 285, 34885–34898 (2010).

12. Subedi, S., Sasidharan, S., Nag, N., Saudagar, P. & Tripathi, T. Amyloid Cross-Seeding: Mechanism, Implication, and Inhibition. Molecules 27, 1776 (2022).

13. Frost, B. & Diamond, M. I. Prion-like mechanisms in neurodegenerative diseases. Nat. Rev. Neurosci. 2009 113 **11**, 155–159 (2009).

14. Luk, K. C. et al. Intracerebral inoculation of pathological α-synuclein initiates a rapidly progressive neurodegenerative α-synucleinopathy in mice. J. Exp. Med. 209, 975–986 (2012).

15. Masuda-Suzukake, M. et al. Prion-like spreading of pathological α-synuclein in brain. Brain 136, 1128–1138 (2013).

16. Goedert, M. Alzheimer’s and Parkinson’s diseases: The prion concept in relation to assembled Aβ, tau, and α-synuclein. Science 349, 1255555 (2015).

17. Goedert, M., Masuda-Suzukake, M. & Falcon, B. Like prions: the propagation of aggregated tau and α-synuclein in neurodegeneration. Brain 140, 266–278 (2017).

18. Soto, C. Unfolding the role of protein misfolding in neurodegenerative diseases. Nat. Rev. Neurosci. 4, 49–60 (2003).

19. Behl, C., Davis, J. B., Lesley, R. & Schubert, D. Hydrogen peroxide mediates amyloid beta protein toxicity. Cell 77, 817–827 (1994).

20. Hsu, L. J. et al. alpha-synuclein promotes mitochondrial deficit and oxidative stress. Am. J. Pathol. 157, 401–410 (2000).

21. Pieri, L., Madiona, K., Bousset, L. & Melki, R. Fibrillar α-Synuclein and Huntingtin Exon 1 Assemblies Are Toxic to the Cells. Biophys. J. 102, 2894–2905 (2012).

22. Mahul-Mellier, A.-L. et al. Fibril growth and seeding capacity play key roles in α-synuclein-mediated apoptotic cell death. Cell Death Differ. 22, 2107–2122 (2015).

23. Eisele, Y. S. et al. Targeting protein aggregation for the treatment of degenerative diseases. Nat. Rev. Drug Discov. 14, 759–780 (2015).

24. Kumar, V. et al. Protein aggregation and neurodegenerative diseases: From theory to therapy. Eur. J. Med. Chem. 124, 1105–1120 (2016).

25. Bourdenx, M. et al. Protein aggregation and neurodegeneration in prototypical neurodegenerative diseases: Examples of amyloidopathies, tauopathies and synucleinopathies. Prog. Neurobiol. 155, 171–193 (2017).

26. Sweeney, P. et al. Protein misfolding in neurodegenerative diseases: implications and strategies. Transl. Neurodegener. 2017 61 **6**, 1–13 (2017).

27. Arar, S., Haque, M. A. & Kayed, R. Protein aggregation and neurodegenerative disease: Structural outlook for the novel therapeutics. Proteins Struct. Funct. Bioinforma. (2023) doi:10.1002/PROT.26561.

28. Galkin, M., Priss, A., Kyriukha, Y. & Shvadchak, V. Navigating α-Synuclein Aggregation Inhibition: Methods, Mechanisms, and Molecular Targets. Chem. Rec. 24, e202300282 (2024).

29. Scheres, S. H., Zhang, W., Falcon, B. & Goedert, M. Cryo-EM structures of tau filaments. Curr. Opin. Struct. Biol. 64, 17–25 (2020).

30. Kumar, S. T. et al. Seeding the aggregation of TDP-43 requires post-fibrillization proteolytic cleavage. Nat. Neurosci. 2023 266 **26**, 983–996 (2023).

31. Mishra, S. Emerging Trends in Cryo-EM-based Structural Studies of Neuropathological Amyloids. J. Mol. Biol. 435, 168361 (2023).

32. Scheres, S. H. W., Ryskeldi-Falcon, B. & Goedert, M. Molecular pathology of neurodegenerative diseases by cryo-EM of amyloids. Nature 621, 701–710 (2023).

33. Todd, T. W., Islam, N. N., Cook, C. N., Caulfield, T. R. & Petrucelli, L. Cryo-EM structures of pathogenic fibrils and their impact on neurodegenerative disease research. Neuron 112, 2269–2288 (2024).

34. Guerrero-Ferreira, R., Kovacik, L., Ni, D. & Stahlberg, H. New insights on the structure of alpha-synuclein fibrils using cryo-electron microscopy. Curr. Opin. Neurobiol. 61, 89–95 (2020).

35. Li, B. et al. Cryo-EM of full-length α-synuclein reveals fibril polymorphs with a common structural kernel. Nat. Commun. 9, 3609 (2018).

36. Guerrero-Ferreira, R. et al. Two new polymorphic structures of human full-length alpha-synuclein fibrils solved by cryo-electron microscopy. eLife 8, e48907 (2019).

37. Schweighauser, M. et al. Structures of α-synuclein filaments from multiple system atrophy. Nature 585, 464–469 (2020).

38. Matiiv, A. B. et al. Structure and Polymorphism of Amyloid and Amyloid-Like Aggregates. Biochem. Mosc. 87, 450–463 (2022).

39. Fitzpatrick, A. W. P. et al. Cryo-EM structures of tau filaments from Alzheimer’s disease. Nature 547, 185–190 (2017).

40. Shi, Y. et al. Cryo-EM structures of tau filaments from Alzheimer’s disease with PET ligand APN-1607. Acta Neuropathol. (Berl.) 141, 697–708 (2021).

41. Falcon, B. et al. Novel tau filament fold in chronic traumatic encephalopathy encloses hydrophobic molecules. Nat. 2019 5687752 **568**, 420–423 (2019).

42. Falcon, B. et al. Structures of filaments from Pick’s disease reveal a novel tau protein fold. Nat. 2018 5617721 **561**, 137–140 (2018).

43. Yang, Y. et al. Structures of α-synuclein filaments from human brains with Lewy pathology. Nature 610, 791–795 (2022).

44. Yang, Y. et al. New SNCA mutation and structures of α-synuclein filaments from juvenile-onset synucleinopathy. Acta Neuropathol. (Berl.) 145, 561–572 (2023).

45. Zhao, K., et al. Parkinson’s disease-related phosphorylation at Tyr39 rearranges α-synuclein amyloid fibril structure revealed by cryo-EM. Proc. Natl. Acad. Sci. U. S. A. 117, 20305–20315 (2020).

46. Hu, J. et al. Phosphorylation and O-GlcNAcylation at the same α-synuclein site generate distinct fibril structures. Nat. Commun. 15, 2677 (2024).

47. Li, D. & Liu, C. Molecular rules governing the structural polymorphism of amyloid fibrils in neurodegenerative diseases. Structure 31, 1335–1347 (2023).

48. Lövestam, S. et al. Seeded assembly in vitro does not replicate the structures of α-synuclein filaments from multiple system atrophy. FEBS Open Bio 11, 999–1013 (2021).

49. Lövestam, S. et al. Assembly of recombinant tau into filaments identical to those of Alzheimer’s disease and chronic traumatic encephalopathy. eLife 11, e76494 (2022).

50. Kudo, Y. Development of amyloid imaging PET probes for an early diagnosis of Alzheimer’s disease. Minim. Invasive Ther. Allied Technol. 15, 209–213 (2006).

51. Alam, M. M., Lee, S. H., Wasim, S. & Lee, S.-Y. PET tracer development for imaging α-synucleinopathies. Arch. Pharm. Res. (2025) doi:10.1007/s12272-025-01538-0.

52. Kallinen, A. & Kassiou, M. Tracer development for PET imaging of proteinopathies. Nucl. Med. Biol. 114–115, 115–127 (2022).

53. Kudo, Y. et al. 2-(2-[2-Dimethylaminothiazol-5-yl]ethenyl)-6-(2-[fluoro]ethoxy)benzoxazole: a novel PET agent for in vivo detection of dense amyloid plaques in Alzheimer’s disease patients. J. Nucl. Med. Off. Publ. Soc. Nucl. Med. 48, 553–561 (2007).

54. Klunk, W. E. et al. Imaging brain amyloid in Alzheimer’s disease with Pittsburgh Compound-B. Ann. Neurol. 55, 306–319 (2004).

55. Fodero-Tavoletti, M. T. et al. In vitro characterisation of BF227 binding to α-synuclein/Lewy bodies. Eur. J. Pharmacol. 617, 54–58 (2009).

56. Tao, Y. et al. Structural mechanism for specific binding of chemical compounds to amyloid fibrils. Nat. Chem. Biol. 19, 1235–1245 (2023).

57. Kikuchi, A. et al. In vivo visualization of α-synuclein deposition by carbon-11-labelled 2-[2-(2-dimethylaminothiazol-5-yl)ethenyl]-6-[2-(fluoro)ethoxy]benzoxazole positron emission tomography in multiple system atrophy. Brain 133, 1772–1778 (2010).

58. Morris, E. et al. Diagnostic accuracy of 18F amyloid PET tracers for the diagnosis of Alzheimer’s disease: a systematic review and meta-analysis. Eur. J. Nucl. Med. Mol. Imaging 43, 374 (2016).

59. Xiang, J. et al. Development of an α-synuclein positron emission tomography tracer for imaging synucleinopathies. Cell 186, 3350–3367.e19 (2023).

60. Liu, K., et al. Binding adaptability of chemical ligands to polymorphic α-synuclein amyloid fibrils. Proc. Natl. Acad. Sci. 121, e2321633121 (2024).

61. Ni, R. & Nitsch, R. M. Recent Developments in Positron Emission Tomography Tracers for Proteinopathies Imaging in Dementia. Front. Aging Neurosci. 13, (2022).

62. Ferrie, J. J. et al. Identification of a nanomolar affinity α-synuclein fibril imaging probe by ultra-high throughput in silico screening. Chem. Sci. 11, 12746–12754 (2020).

63. Chia, S. et al. Structure-Based Discovery of Small-Molecule Inhibitors of the Autocatalytic Proliferation of α-Synuclein Aggregates. Mol. Pharm. 20, 183–193 (2023).

64. Gainza, P. et al. Deciphering interaction fingerprints from protein molecular surfaces using geometric deep learning. Nat. Methods 17, 184–192 (2020).

65. Schmid, A. W., Fauvet, B., Moniatte, M. & Lashuel, H. A. Alpha-synuclein Post-translational Modifications as Potential Biomarkers for Parkinson Disease and Other Synucleinopathies. Mol. Cell. Proteomics 12, 3543–3558 (2013).

66. Altay, M. F. et al. Development and validation of an expanded antibody toolset that captures alpha-synuclein pathological diversity in Lewy body diseases. Npj Park. Dis. 9, 1–21 (2023).

67. Krivák, R. & Hoksza, D. P2Rank: machine learning based tool for rapid and accurate prediction of ligand binding sites from protein structure. J. Cheminformatics 10, 39 (2018).

68. Seidler, P. M. et al. Structure-based discovery of small molecules that disaggregate Alzheimer’s disease tissue derived tau fibrils in vitro. Nat. Commun. 2022 131 13, 1–12 (2022).

69. Cao, T., Li, X., Li, D. & Tao, Y. Development of small molecules for disrupting pathological amyloid aggregation in neurodegenerative diseases. Ageing Neurodegener. Dis. 3, N/A-N/A (2023).

70. Limorenko, G. & Lashuel, H. A. To target Tau pathologies, we must embrace and reconstruct their complexities. Neurobiol. Dis. 161, 105536 (2021).

71. Limorenko, G. & Lashuel, H. A. Revisiting the grammar of Tau aggregation and pathology formation: how new insights from brain pathology are shaping how we study and target Tauopathies. Chem. Soc. Rev. 51, 513–565 (2022).

72. Lashuel, H. A. Rethinking protein aggregation and drug discovery in neurodegenerative diseases: Why we need to embrace complexity? Curr. Opin. Chem. Biol. 64, 67–75 (2021).

73. Khare, S. D., Chinchilla, P. & Baum, J. Multifaceted Interactions Mediated by Intrinsically Disordered Regions Play Key Roles in Alpha Synuclein Aggregation. Curr. Opin. Struct. Biol. 80, 102579 (2023).

74. Donzelli, S. et al. Post-fibrillization nitration of alpha-synuclein abolishes its seeding activity and pathology formation in primary neurons and in vivo. 2023.03.24.534149 Preprint at 10.1101/2023.03.24.534149 (2023).

75. Balana, A. T. et al. O-GlcNAc forces an α-synuclein amyloid strain with notably diminished seeding and pathology. Nat. Chem. Biol. 20, 646–655 (2024).

76. Horne, R. I. et al. Discovery of potent inhibitors of α-synuclein aggregation using structure-based iterative learning. Nat. Chem. Biol. 20, 634–645 (2024).

77. Word, J. M., Lovell, S. C., Richardson, J. S. & Richardson, D. C. Asparagine and glutamine: using hydrogen atom contacts in the choice of side-chain amide orientation. J. Mol. Biol. 285, 1735–1747 (1999).

78. Sanner, M. F., Olson, A. J., Jolla, L. & Spehner, J.-C. Reduced Surface: An Efficient Way to Compute Molecular Surfaces. J. Mol. Graph. 11, 139–1411 (1993).

79. Zhou, Q. PyMesh—Geometry Processing Library for Python. Software available for download at. (2019).

80. Baker, N. A., Sept, D., Joseph, S., Holst, M. J. & McCammon, J. A. Electrostatics of nanosystems: Application to microtubules and the ribosome. Proc. Natl. Acad. Sci. U. S. A. 98, 10037–10041 (2001).

81. Kyte, J. & Doolittle, R. F. A simple method for displaying the hydropathic character of a protein. J. Mol. Biol. 157, 105–132 (1982).

82. Yin, S., Proctor, E. A., Lugovskoy, A. A. & Dokholyan, N. V. Fast screening of protein surfaces using geometric invariant fingerprints. Proc. Natl. Acad. Sci. U. S. A. 106, 16622–16626 (2009).

83. Kortemme, T., Morozov, A. V. & Baker, D. An Orientation-dependent Hydrogen Bonding Potential Improves Prediction of Specificity and Structure for Proteins and Protein–Protein Complexes. J. Mol. Biol. 326, 1239–1259 (2003).

84. Zhou, Q.-Y., Park, J. & Koltun, V. Open3D: A Modern Library for 3D Data Processing. Preprint at 10.48550/arXiv.1801.09847 (2018).

85. Cock, P. J. A. et al. Biopython: freely available Python tools for computational molecular biology and bioinformatics. Bioinformatics 25, 1422–1423 (2009).

86. Hallinan, G. I. et al. Cryo-EM structures of prion protein filaments from Gerstmann– Sträussler–Scheinker disease. Acta Neuropathol. (Berl.) 144, 509–520 (2022).

87. Yang, Y. et al. Cryo-EM structures of amyloid-b 42 filaments from human brains. Science 375, 167–172 (2022).

88. Schweighauser, M. et al. Age-dependent formation of TMEM106B amyloid filaments in human brains. Nat. 2022 6057909 605, 310–314 (2022).

89. Boyer, D. R. et al. The α-synuclein hereditary mutation E46K unlocks a more stable, pathogenic fibril structure. Proc. Natl. Acad. Sci. 117, 3592–3602 (2020).

90. Boyer, D. R. et al. Structures of fibrils formed by α-synuclein hereditary disease mutant H50Q reveal new polymorphs. Nat. Struct. Mol. Biol. 26, 1044–1052 (2019).

91. Sun, Y. et al. The hereditary mutation G51D unlocks a distinct fibril strain transmissible to wild-type α-synuclein. Nat. Commun. 2021 121 12, 1–10 (2021).

92. Sun, Y. et al. Cryo-EM structure of full-length α-synuclein amyloid fibril with Parkinson’s disease familial A53T mutation. Cell Res. 2020 304 30, 360–362 (2020).

